# Sugar Beet Extract Acts as a Natural Bio-Stimulant for Physio-Biochemical analysis of *Hordeum Vulgare* L. under Induced Salinity Stress

**DOI:** 10.1101/2021.08.19.457015

**Authors:** Noor Ali Shah, Sami Ullah, Muhammad Nauman Khan, Sajjad Ali, Muhammad Adnan, Amjad Ali, Muhammad Ali, Ajmal Khan, Said Hassan, Wisal Muhammad Khan

## Abstract

Change in climate of the entire globe due to elevated temperature and minimum annual rainfall in barren zone frequently leads to salinity of soil. The current study was aimed to evaluate the importance of sugar beet extract (SBE) as a bio-stimulant to improve the adverse damages of induced salinity stress (40mM) on growth, oosmolytes and antioxidant defense system of barley (*Hordeum vulgare* L.). Pot experiment was carried in green house under different concentrations of SBE (10%, 20%, 30%, 40%, 50%) pre-soaked seeds of *Hordeum vulgare* for 5 hours SBE was analyzed for glycine betaine (100mmol/kg), betalains (1.3mg/l), phenolics (1.30g/100ml), flavonoids (0.59mg/ml), carotenoids (0.23ml/100ml), vitamin E (0.002%), vitamin C (8.04g/100ml), sugar (8g/100ml), protein (1.39mg/100ml), and oxalic acid (38mg/100ml) while Ca (13.72mg/l), Mg (7.121 mg/l) and K (11.45mg/l) contents were also determined. We found significant improvement in germination parameters of *Hordeum vulgare* L. via SB extract on coefficient of velocity of emergence (CVE), mean emergence time (MET), germination energy (GE), timson germination index (TGI), germination rate index (GRI) and time to 50% emergence (E_50_) under induced salinity stress. However, photosynthetic pigments, e.g., chlorophyll and carotenoids were enhanced using 40% SB extract, soluble sugar, protein, proline, POD, MDA with 50% SB extract while SOD and H_2_O_2_ in 20% SBE, respectively. Our findings suggested that SB extract promotes both agronomical and physiological attributes, is a positive way to enhance our economy by increasing crop yields in arid and semi-arid areas along with plant tolerance to under induced salinity stress.

**Highlights:** The recent study is of utmost importance owing to the following reasons:

➢ Changes in climatic condition results in salinity, droughts, floods, earthquakes, fluctuation in temperature and other environmental hazards.
➢ Salinity stress has myriad of damaging effects on crop growth and productivity and becoming a global issue of major concern especially in the Asian countries including Pakistan. Statistical analysis showed that 6.30 million of lands is affected with salinity.
➢ Barley (*Hordeum Vulgare* L.) being an important crop of economic and nutritional value has been facing salinity stress condition in Pakistan, resulting in productivity reduction and ultimately paving the way for food scarcity and crippling economy.
➢ The sole purpose of this research article is to assess and ameliorate the physio-biochemical responses via application of sugar beet extract as bio-stimulants in Barley (*Hordeum Vulgare* L.) grown under induced salinity stress; in order to cope with the dire consequences of salinity stress especially in the southern areas of Pakistan being badly affected by salt stress.
➢ Furthermore, there is a great need for adaptive measures suggested by the present research and other findings made by researchers in pursuit of tackling this alarming global menace.

## 1. Introduction

By the end of 2050, abiotic stresses such as salinity, drought and temperature will lead to huge losses in crop and food production (Ahmed et al. 2020; Hawrylak et al. 2019). The beginning of the 21st century is marked by global shortage of water and soil salinity (Sanders 2020; Fedoroff et al. 2010). Salinity is one of the main environmental factors which limiting plants productivity (Munns and Tester 2008). Worldwide, salinity has been reported that about 7.0% (which includes an estimated area of about 77 Mha) of the world’s land surface is occupied by various soils affected by salinity (Kashif et al. 2020). Salinity causes many abnormal changes in the morphological and physiological functioning of plant cells due to sodium and chloride ions in cells, disturbs the level of minerals and reduces the water potential of the soil for sowing by a high osmotic potential causing high osmotic stress (Zhu et al 2020). Due to salinity, synthesis of lipids and protein along with photosynthesis are badly affected in plants (Javed et al. 2020). To respond to such reactive oxygen species (ROS), plants immediately regulate their antioxidant defense mechanism by activating antioxidant enzymes activities; such as SOD, POD, CAT (Amirjani 2011). Catalase enzyme has a greater importance to convert H_2_O_2_ into useful water and oxygen, while peroxidases break down H_2_O_2_ through the oxidation of the cosubstrate which includes phenolic compounds and other antioxidants. Similarly, the ascorbate peroxidase enzyme synthesizes ascorbic acid (AsA) as an electron donor in the antioxidant process that increases H_2_O_2_ (Prasad et al. 2009).

Barley belongs to the family Poaceae and is the 4^th^ most important cereal crop (Gorzolka et al. 2016). It is the 4^th^ most important and stress tolerant cereal crop globally, after wheat, maize and rice (Imran et al. 2020). About 6.30 million of lands is affected with salinity (Economic survey of Pakistan 2017). According to an estimate about 20% of all irrigated and cultivated lands (equivalent to 62 million ha) are negatively affected by salt stress at present time (Javed et al. 2020). Production of barley fluctuated substantially in the history of Pakistan. It occupies an area of 84.1 thousand hectares with a production of 71.4 thousand tons (Ali et al. 2020). Being a simple and cost effective, seed priming technique has been used to improve plant growth, germination and enhanced seedling’s vigor (Zhang and Rue 2014). Application of SBE and GB on the *Solanum melongena* L. plants to alleviate salinity stress. Since sugar beet extract contains important nutrients like AsA, GB, vitamin E, sugars and amino acids, etc. Also, it can be used as a cheaper source of GB to improve and protect plants from the lethal effect of salt stress (Abbas et al. 2010; Ahanger et al. 2020). All the compounds in sugar beet, that is, play a vital role as a bio-stimulant in plant life during stressful conditions. Seed priming technique, best pregermination seed treatment has been revealed to boost seedling and yield germination (Nasri et al. 2013). The application of SBE as bio-stimulants is boosting speedily in agronomic crops for valuable effects under stressful conditions (Pinheiro et al. 2018; Mahdy et al. 2020). It was assumed priming of seeds with SBE could lower the damaging effects of salinity stress on seed seedling emergence, germination, and establishment. Therefore, the present experiment was performed to evaluate the capability of seed pre-sowing with SBE to regulate barley seed germination under 40 mM NaCl stress. In addition, to explore the degree of efficacy of SBE responses in terms of physiological and biochemical attributes were kept in focus to find out a viable strategy for blocking oxidative stress in barley.

## 2. Material and Methods

### 2.1. Formation of Sugar beet extract (SBE)

The fresh roots of *Beta vulgaris* L. were obtained from local market, grind to obtain juice and stored at 10°C. Different concentrations of SBE i.e. (10%, 20%, 30%, 40% and 50% v/v) were then prepared and applied for the treatments.

### 2.2. Estimation of glycine betaine (GB) and Betalain content

SB extract was analyzed for glycine betaine (GB) by following the methodology of Grieve and Grattan (1983). Betalain content was investigated by the following method Nilsson (1970), the absorbance of betalains were measured at 538 and 480 nm using the method of Cai (2005) equation.

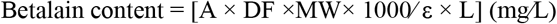

DF = dilute factor, MW = molecular weight, and ε is the extinction coefficient, for indicaxanthin and betanin, i.e. 48,000 and 60,000 1/mol.

### 2.3. Estimation of phenolic and flavonoid content

Sugar beet extract was examined phenolics in SBE following method of Waterhouse (2002) with optical density at 765 nm. Similarly, flavonoid content was calculated as per described by the methodology of Bushra et al. (2009) by reacting the dilute SBE with 0.5% of NaNO_2_ and 10% AlCl_3_. After that, 0.2 ml of sodium hydroxide was added. optical density was recorded at 415 nm.

### 2.4. Estimation of carotenoid content

Carotenoid contents in sugar beet extract were determined by Mohdaly et al. (2010). Using spectrophotometer and measured at 450 nm by the following equation.

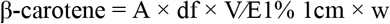

Here, df = dilution factor, V = volume (ml), E1% = coefficient of absorbency for petroleum ether was (2592)1 cm, w = weight of sample (g), A = absorbance.

### 2.5. Estimation of alpha-tocopherol (Vitamin E) and ascorbic acid (Vitamin C)

According to Backer et al. (1980) method was used to study alpha-tocopherol content in sugar beet extract at 520nm whereas, following the standard protocol of *Mukherjee* and *Choudhuri* (1983) ascorbic acid was determined in sugar beet extract. The absorbance of vitamin C was measured at 530 nm.

### 2.6. Estimation of total soluble sugars (TSS) and oxalic acid content

Total soluble sugars were investigated by using the method of Dubois et al. (1956). Similarly, oxalic acid content in SBE was estimated by per manganese method. Oxalic acid concentrations were determined using a curve generated by an oxalic acid.

### 2.7. Estimation of SBE minerals and pH

Flame photometer was used for the estimation of Ca, P, K, and Mg contents, while Atomic Absorption Spectrometry (AAS700, Perkin Elmer, USA), and was applied to determine Mn, Fe, Cu and Ni, contents in SBE, Centralized Resource Laboratory (CRL), University of Peshawar, Pakistan. However, pH of the sugar beet extract measured with pH meter (WTW Inolab 7110).

### 2.8. Site description and experimental design

Field experiment was carried out at Department of Botany, University of Peshawar (34° 1’ 33.3012” N and 71° 33’ 36.4860’’ E.) during barley growing season in 2018. Barley seeds (*Hordeum vulgare* L.) were collected from National Agricultural Research Center (NARC), Islamabad, Pakistan. Intact seeds of uniform shape and size were surface sterilized 0.3% sodium hypochlorite for three minutes and the rinsed and soaked in sugar beet extract with different concentrations of SBE such as, 10%, 20%, 30%, 40% and 50% for 5 hours. The seeds were sown in earthen pots (20 cm height), with upper/lower diameter (18cm) filled with 4 kg of sandy and loamy (2:1) having pH (6.0), EC (2.41ds/m), Organic Carbon (C) 22.6 g kg^-1^ (Nelson and Sommers 1983), Nitrogen (N) content 3.09 g kg^-1^ (Keeney and Nelson, 1983), Potassium (K) available 92.3 mg kg^-1^ (Nelson and Heidel 1952) and Phosphorus (P) 8.0 mg kg^-1^ (Jacson et al. 1973) and percent moisture content of 46%. Each pot was kept at 15cm apart in a complete randomized design and were protected from rain. About 10 seeds of barley have been sown in each pot. Experiment was divided into 11 treatments (T1= Control (untreated), T2= 10 % SBE, T3= 10 % SBE + 40 mM NaCl, T4= 20% SBE, T5= 20% SBE + 40 mM NaCl, T6= 30% SBE, T7=30% SBE + 40 mM NaCl, T8= 40% SBE, T9= 40% SBE + 40 mM NaCl, T10=50% SBE, T11= 50% SBE + 40 mM NaCl) with three replicates per treatment. Hand weeding, watering and plant thinning were done, and the seedlings were exposed to proper sunlight for healthier growth. Salinity stress (40 mM NaCl) was applied after 25 days of post germination, while plant samples for the determination of growth and physiological parameters were collect after 10 days of saline treatment and preserved in refrigerator at 4°C.

### 2.9. Elemental analysis of plant foliar material

EDX technique (INCA100/ Oxford instruments, U.K.) was performed for elemental analysis in barley leaves including carbon (C), oxygen (O), nitrogen (N), Sulphur (S), Phosphorus (P), Sodium (Na), Calcium (Ca), Magnesium (Mg), Potassium (K), Chlorine (Cl) and Silicon (Si).

### 2.10. Agronomic characteristics

#### 2.10.1. Mean Germination Time (MGT)

Mean germination time for seed emergence investigated by Kader (2005). Greater will be the inhabitants germinated lower is mean germination time.

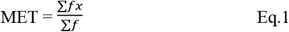

Hence “f” shows the numbers of seed germinated on X days.

#### 2.10.2. Time to 50% Germination (T50%)

Time to 50% germination planned by the Vujosevic et al. (2018).

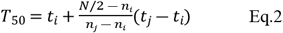

Hence “N” refers to the final emerged seeds number, where nj and ni are inconsistent number of seeds germinated by the adjoining count at tj and ti, correspondingly, when ni<N/2>Nj.

#### 2.10.3. Germination Rate Index (GRI)

Germination rate index was calculated by Kaydan and Yagmur (2008).

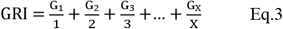

Hence “G1” is the germinated seed percentage at day first and “G2” is germinated seed percentage at second day after sowing.

#### 2.10.4. Coefficient of Velocity of Emergence (CVE)

Coefficient of velocity of emergence was determined by the method of Kader (2005).

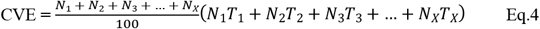

Hence “n” is the number of seed germinated per/day while T represents the time.

#### 2.10.5. Timson Germination Index (TGI), Germination Energy (GE)

Timson germination index and Germination energy was measured by the method of Al-Ansari and Siksi (2016).

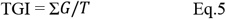

Hence G is the germination percentage per day, while T is the entire germination period.

Germination energy was determined by the following formula Al-Ansari and Siksi (2016).

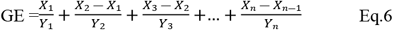

Hence, X_n_ = numbers of seed germination on last (nth) counting date and Y_n_ represents the number of days from sowing to last (nth) counting date.

### 2.11. Physiological and biochemical analysis of plant foliar material

#### 2.11.1. Determination of Chlorophyll and Carotenoid Content

Chlorophyll and carotenoid contents were determined by the method of Marcinska et al. (2012) with some modifications. Foliar material of 0.2 g was chopped in 80% acetone and incubated for 24 hours in dark then centrifuged. Absorbance values for chlorophyll “a” at 649 nm, chlorophyll “b at 663 nm and carotenoid at 470 nm were observed against 80% acetone blank through spectrophotometer.

#### 2.11.2. Estimation of Osmolytes Content

Leaf soluble sugar content was evaluated by the methodology of Dubois et al. (1956). Foliar material of 0.5 g was grounded in 5 ml distilled water and centrifuged for 10 minutes. About 4 ml of 35% concentrated H_2_SO_4_ was added to 1ml supernatant its optical density was found at 490 nm.

Total proline content was measured by the methodology of Plazek et al. (2013). 0.5 g of fresh leaves were grounded 3% (10 ml) sulfosalicylic acid and filtered. After filtration, 2 ml of the filtrate was dissolved in 2 ml acid ninhydrin (40 ml glacial acetic acid + 1.87 g ninhydrin + 30 ml of 6 M phosphoric acid) and glacial acetic acid in a test tube and warm for 1 h at 100°C. Solution extraction was completed with toluene (4 ml) and OD was measured at 520 nm.

#### 2.11.3. Estimation of Soluble Protein (SP) and Hydrogen Peroxide Content (H_2_O_2_)

Soluble protein was found by the standard protocol of Rostami and Ehsanpaur (2009) with some modifications. 0.5 g foliar material was grounded in 1 ml phosphate buffer solution (pH 7.0). After homogenation for 10 min, about 0.1 ml extract mixed 0.9 ml distilled water and 1 ml reagent in 40 ml distilled water (0.1 N sodium hydroxide, 0.75 g sodium carbonate, 0.37 g sodium potassium tartrate). By shaking and adding 0.1 ml foline phenol was kept for incubation of 30 min and the absorbance was measured at 650 nm over spectrophotometer.

H_2_O_2_ concentration was investigated by *Velikova* et al. (2000). Fresh foliar material (0.5 gm) was chopped in 5 ml trichloro acetic acid and centrifuged for 10 min. About 0.5 ml of the supernatant was dissolved in 0.5 ml phosphate buffer and 1 ml potassium iodide reagent. Optical density was determined at 390 nm.

#### 2.11.4. Determination of Antioxidant Enzymes

Leaf MDA content was estimated by Zhang and Huang (2013). 0.25 g of leaves were homogenized in 3 ml of 1.0% Trichloro acetic acid and centrifuge for 10 minutes. 1 ml of the supernatant was mixed with 0.5% (4 ml) 2-thiobarbituric acid and heated for one hour at 95 °C, then cooled with ice for 10 minutes and its optical density was determined at 532 nm.

SOD activity was determined by the method of Wang et al. (2014) with a little bit change. 3 ml reaction mixture contained 0.1 ml supernatant, 0.72 ml methionine (58 mg), 0.72 ml nitroblue tetrazolium (1.89 mg), 0.72 ml ethylene diamine tetra acetic acid (1.1 mg) and 0.72 ml riboflavin (0.02 mg) in 120 ml distilled water and then kept for incubation in dark and light period of 30 min. Optical density was determined by spectrophotometer at 560 nm.

POD activity of plant foliar material was performed by the methodology of Asthir et al. (2009). 0.5 g fresh leaves were chopped with 2 ml solution comprises of 12.5 g polyvinyl pyrrolidine, 4.6 g ethylene diamine tetra acetic acid, 0.2 ml phosphate buffer solution with 7.0 pH in 125 ml of distilled water and then centrifuged for 20 min. 3 ml reaction mixture contained supernatant (0.1 ml), methyl ethyl sulfonic acid buffer (1.3 ml), phenyl diamine (0.1 ml) and a drop of H_2_O_2_ (3%). Optical density was measured at 485 nm by spectrophotometer.

### 2.12. Statistical Analysis

The statistical analysis was three factors (barley, SBE, salinity) which arranged in CRD. Data collected for various germination attributes, physiological and biochemical components by using IBM SPSS Statistics 22. Mean separation and standard deviation were determined by Tukey’s test.

## 3. Results

The preliminary work was planned to determine the role of SBE as bio-stimulants under different percent concentrations. Various bio-chemical compounds were investigated in SBE included glycine betaine (100mmol/kg), betalains (1.3mg/l), phenolics (1.30g/100ml), flavonoids (0.59mg/ml), carotenoids (0.23ml/100ml), vitamin E (0.002%), vitamin C (8.04g/100ml), sugar (8g/100ml), protein (1.39mg/100ml), and oxalic acid, while the concentrations of various nutrient minerals such as Ca (13.72mg/l), Mg (7.121 mg/l), K (11.45mg/l), Na (3.098 mg/l), Fe (1.501 mg/l), Cu (0.110 mg/l), Pb (0.005 mg/l), Cd (0.001 mg/l) and Zn (0.191 mg/l) contents were also determined as shown in ‘Table 1’. However elemental analysis of plant foliar material results indicated in ‘Table 2’ included C (58.0-47.5%), O (31.3-39.0%), N (12.7-6.5%), P (0.33-0.5%), S (0.32-0.15%), Na (0.5-0.2%), Ca (0.74-0.25%), Mg (0.18-0.08%), K (2.36-0.08%), Cl (1.02-0.49%) and Si (0.32-0.13%).

**Table 1.** Composition of SBE used as priming seed treatment under salinity stress.

| Biochemical/Nutrients | Concentration |
| --- | --- |
| Glycine betaine | 100 mmol/kg f.wt |
| Betalains | 1.36 mg/l |
| Phenolics | 1.30 g/100ml |
| Flavonoids | 0.59 mg/ml |
| Carotenoids | 0.23 ml/100ml |
| Vitamin E | 0.002% |
| Vitamin C | 8.04g/100ml |
| Sugar | 8 g/100ml |
| Protein | 1.39 mg/100ml |
| Oxalic acid | 38 mg/100ml |
| pH | 6.07 |
| Cu | 0.110 mg/L |
| Pb | 0.005 mg/L |
| Cd | 0.001 mg/L |
| Zn | 0.191mg/L |
| Na | 3.098 mg/L |
| Ca | 13.72 mg/L |
| K | 11.45 mg/L |
| Mg | 7.121 mg/L |
| Fe | 1.501 mg/L |

**Table 2.**
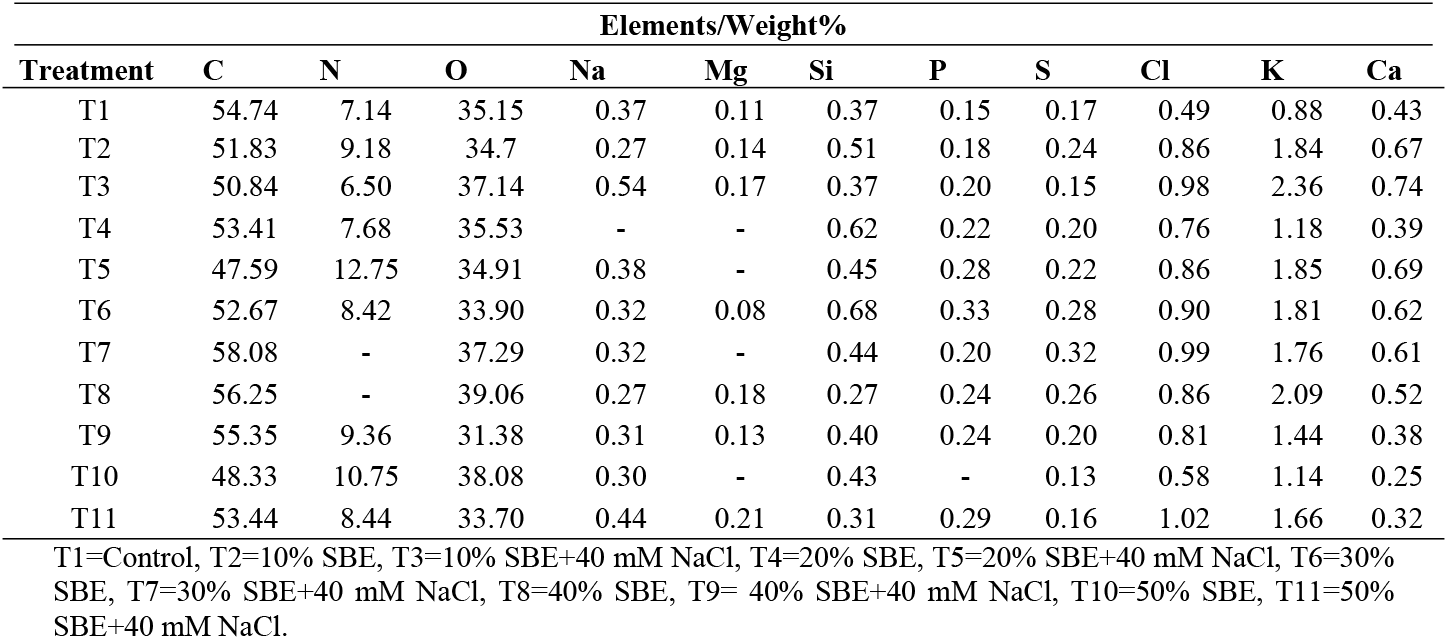
Elemental analysis of plant tissue nutrients is analyzed by EDX under salinity stress.

| Treatment | Elements/Weight% |  |  |  |  |  |  |  |  |  |  |
| --- | --- | --- | --- | --- | --- | --- | --- | --- | --- | --- | --- |
|  | C | N | O | Na | Mg | Si | P | S | Cl | K | Ca |
| T1 | 54.74 | 7.14 | 35.15 | 0.37 | 0.11 | 0.37 | 0.15 | 0.17 | 0.49 | 0.88 | 0.43 |
| T2 | 51.83 | 9.18 | 34.7 | 0.27 | 0.14 | 0.51 | 0.18 | 0.24 | 0.86 | 1.84 | 0.67 |
| T3 | 50.84 | 6.50 | 37.14 | 0.54 | 0.17 | 0.37 | 0.20 | 0.15 | 0.98 | 2.36 | 0.74 |
| T4 | 53.41 | 7.68 | 35.53 | - | - | 0.62 | 0.22 | 0.20 | 0.76 | 1.18 | 0.39 |
| T5 | 47.59 | 12.75 | 34.91 | 0.38 | - | 0.45 | 0.28 | 0.22 | 0.86 | 1.85 | 0.69 |
| T6 | 52.67 | 8.42 | 33.90 | 0.32 | 0.08 | 0.68 | 0.33 | 0.28 | 0.90 | 1.81 | 0.62 |
| T7 | 58.08 | - | 37.29 | 0.32 | - | 0.44 | 0.20 | 0.32 | 0.99 | 1.76 | 0.61 |
| T8 | 56.25 | - | 39.06 | 0.27 | 0.18 | 0.27 | 0.24 | 0.26 | 0.86 | 2.09 | 0.52 |
| T9 | 55.35 | 9.36 | 31.38 | 0.31 | 0.13 | 0.40 | 0.24 | 0.20 | 0.81 | 1.44 | 0.38 |
| T10 | 48.33 | 10.75 | 38.08 | 0.30 | - | 0.43 | - | 0.13 | 0.58 | 1.14 | 0.25 |
| T11 | 53.44 | 8.44 | 33.70 | 0.44 | 0.21 | 0.31 | 0.29 | 0.16 | 1.02 | 1.66 | 0.32 |

T1=Control, T2=10% SBE, T3=10% SBE+40 mM NaCl, T4=20% SBE, T5=20% SBE+40 mM NaCl, T6=30% SBE, T7=30% SBE+40 mM NaCl, T8=40% SBE, T9= 40% SBE+40 mM NaCl, T10=50% SBE, T11=50% SBE+40 mM NaCl.

### 3.1. Germination attributes of *Hordeum vulgare* L

In the present study, the signs of germination showed a significant role of SBE in terms of resistance to salinity stress and maintaining good germination. The results (Table 3) indicated that TGI and CVE enhanced by pre-treating seed of barley with 30% SBE under induced salinity stress while decrease in SBE alone without salinity stress. Similarly, GRI and T_50%_ indicated maximum gemination under 30% SBE treatment whereas reduced in control condition. Consequently, GE and MET have been determined highest in 10% and 40% SBE treatment in the absence of salinity while decrease in 30% SBE pre-treated seeds of barley. On the whole, the results confirmed that concentration of 30% and 40% SBE used for the priming was found to be more useful in terms of G% for the seeds exposed to salinity stress.

**Table 3.** Effect of SBE on timson germination index (TGI), germination energy (GE), coefficient of uniformity of emergence (CUE), mean germination time (MGT), germination rate index (GRI) and time to 50% emergence (T50%) under induced salinity stress.

| Treatment | TGI | GE | CVE | MET | GRI | T50% |
| --- | --- | --- | --- | --- | --- | --- |
| T1 | 1.5±0.06 | 11.11±3.14 | 202.47±10.55 | 26.97±0.67 | 6.53±0.15 | 1.61±0.9 |
| T2 | 3.3±0.11 | 26.66±2.44 | 351.94±15.83 | 34.48±0.82 | 9.75±0.36 | 2.09±0.3 |
| T3 | 3.1±0.04 | 22.22±3.14 | 331.32±14.04 | 33.62±0.81 | 9.31±0.36 | 2.38±0.8 |
| T4 | 2.0±0.24 | 20.21±2.11 | 281.52±44.95 | 30.91±2.23 | 8.51±0.17 | 2.40±0.9 |
| T5 | 1.9±0.19 | 17.77±3.28 | 280.10±28.90 | 30.04±1.32 | 8.35±0.76 | 2.16±0.5 |
| T6 | 1.6±0.43 | 15.55±3.14 | 208.34±97.14 | 26.24±5.76 | 6.94±0.73 | 3.73±0.4 |
| T7 | 3.7±0.12 | 37.77±3.14 | 421.33±25.26 | 26.8±0.44 | 11.33±0.47 | 2.97±0.9 |
| T8 | 1.8±0.17 | 20.00±3.44 | 252.35±15.94 | 37.08±0.99 | 8.08±0.65 | 2.66±0.4 |
| T9 | 1.9±0.14 | 22.22±3.14 | 253.59±34.13 | 29.48±0.59 | 8.35±0.58 | 2.23±0.5 |
| T10 | 1.1±0.11 | 24.44±3.14 | 327.36±25.42 | 29.17±2.01 | 9.31±0.44 | 2.42±0.5 |
| T11 | 2.9±0.22 | 26.66±3.44 | 262.36±30.89 | 33.35±1.24 | 8.55±0.84 | 3.25±0.0 |

T1= Control, T2= 10% SBE, T3=10 % SBE+40 mM NaCl, T4=20% SBE, T5=20% SBE+40 mM NaCl, T6=30% SBE, T7=30% SBE+40 mM NaCl, T8=40% SBE, T9=40% SBE+40 mM NaCl, T10=50% SBE, T11=50% SBE+40 mM NaCl.

### 3.2 Physiological and biochemical attributes

It is suggested that the degradation of plant chlorophyll content and increase in cellular respiration rate under stress condition is related with the accumulation of reactive oxygen species (Camejo et al. 2006). Plant foliar material was analyzed for photosynthetic pigments such as chlorophyll a/b ratio, total Chlorophyll and carotenoid content were significantly enhanced by increasing concentration from 20-40% SBE under salinity stress. According to (Fig. 1-3) results revealed that maximum values of chlorophyll a/b ratio, total chlorophyll content and carotenoid contents were recorded in T5, T10, and T6 while minimum results have been obtained by applying the extract without salinity indicated that SBE is a natural bio-stimulant as well as creating plant ability to stress tolerance by activating different physiological processes respectively. Furthermore, the results showed that there is a big difference among all the primed and non-primed treatments under induced salinity stress (40 mM NaCl).

**Figure 1.**
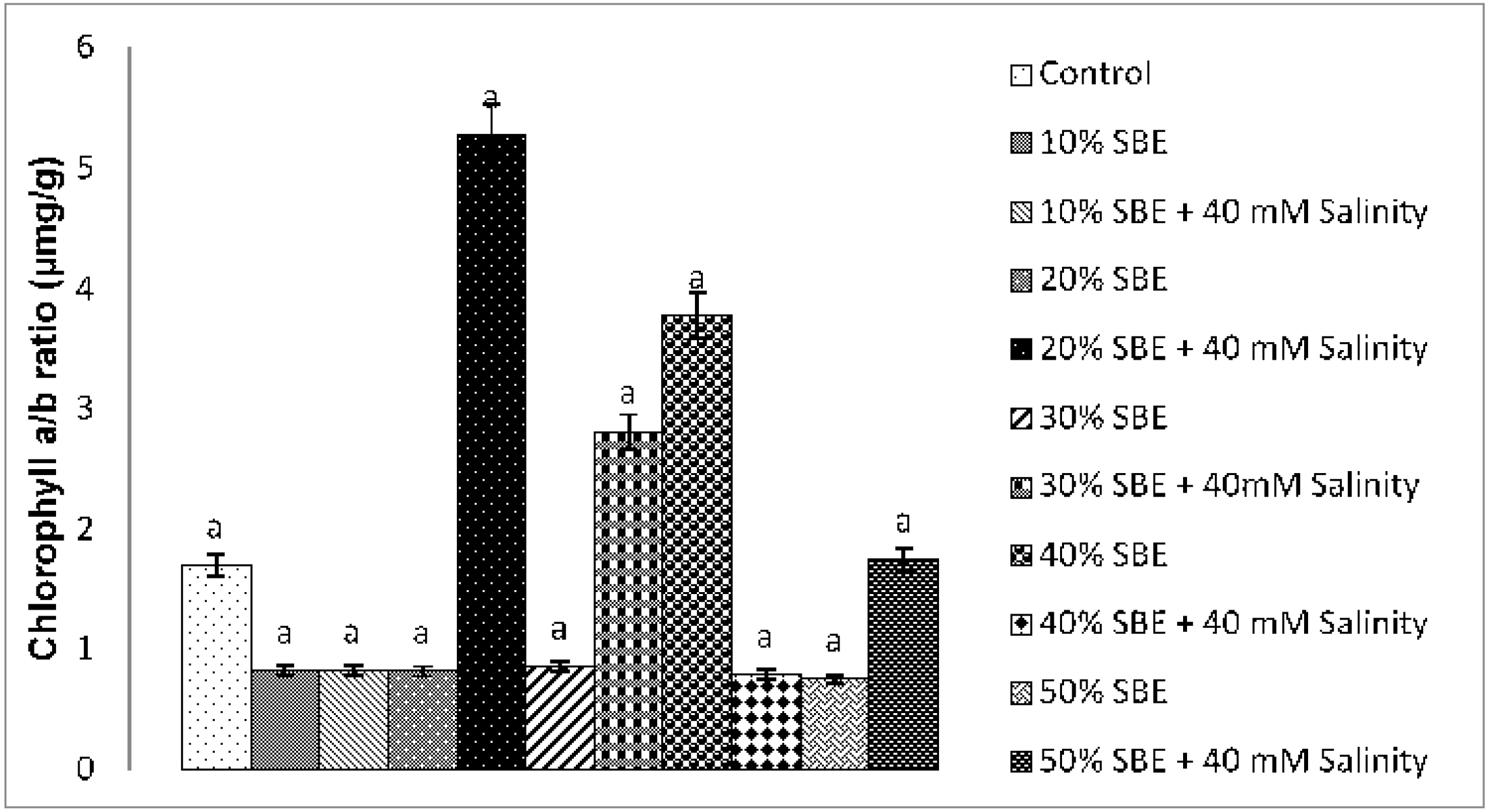
Effect of sugar beet extract on chlorophyll “a/b” ratio under induced salinity stress. Vertical bars represent SE.

**Figure 2.**
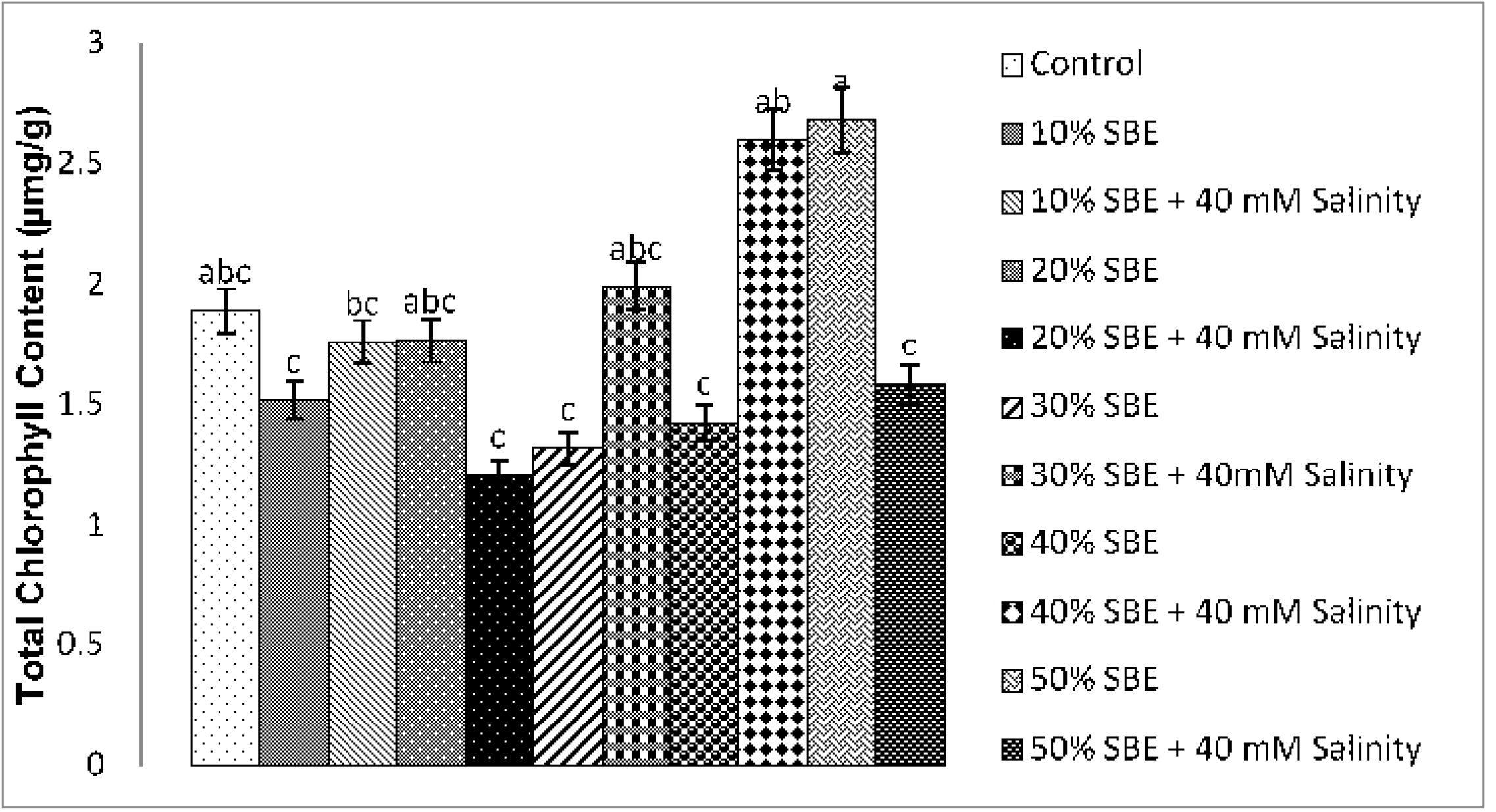
Effect of sugar beet extract on total chlorophyll contents under induced salinity stress. Vertical bars represent SE.

**Figure 3.**
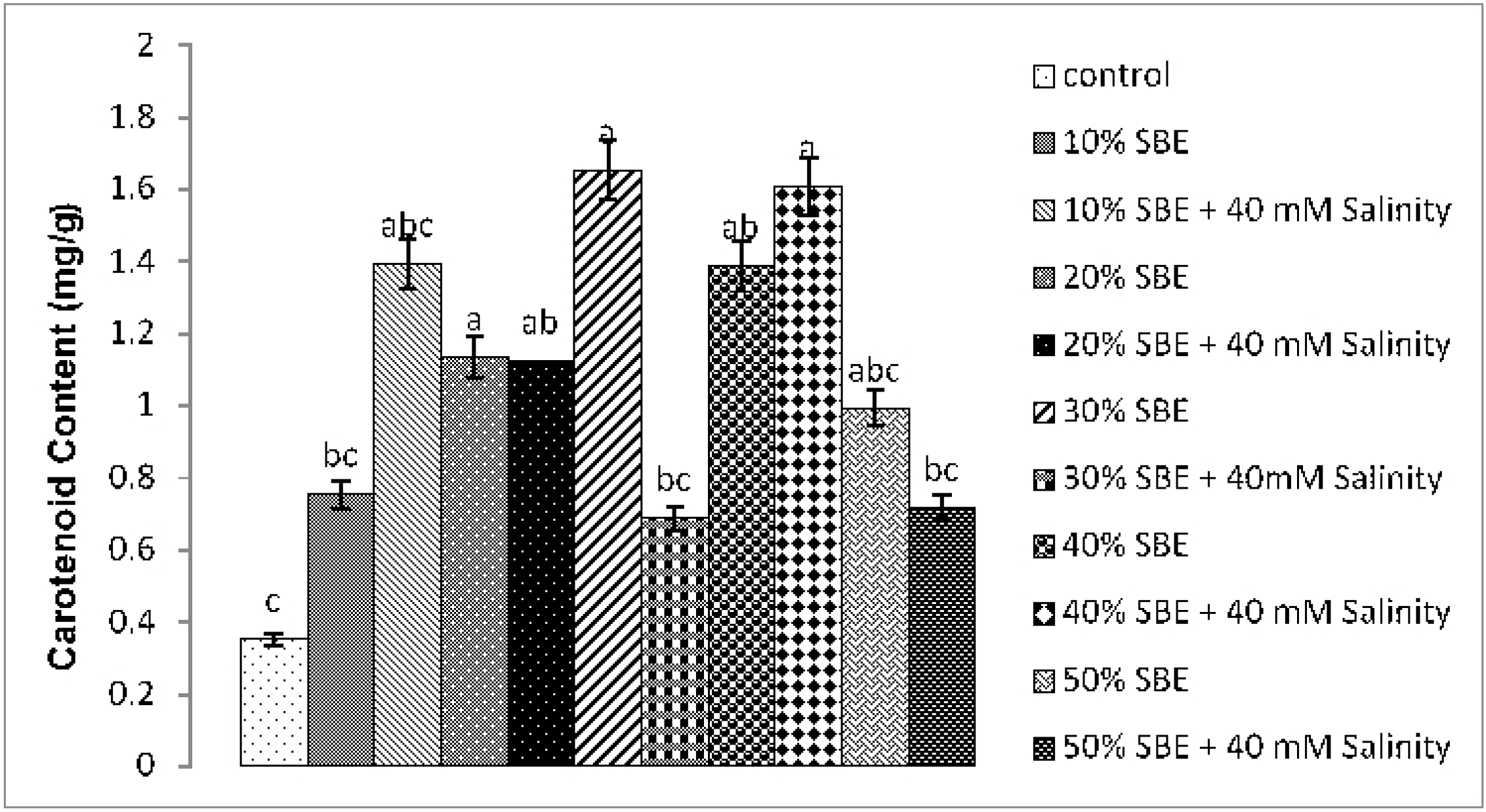
Effect of sugar beet extract on carotenoid content under induced salinity stress. Vertical bars represent SE.

Plants perceive a change in their physiological events in response to several abiotic stress and therefore respond shortly by accumulating different protective osmolytes mostly comprising sugar, proline and glycine betaine to provide favorable environment for metabolic activities and also protecting the plant by reducing the harmful damages caused by oxidative stress (Hayat et al. 2012). Similarly, plant osmolyte such as soluble sugar content has been recorded maximum in T11 followed by T9 under induced salinity stress in (Fig. 4). Results revealed that primed seeds have a great influenced on the soluble sugar content up to a significant level. Therefore, it is investigated that increasing the SBE concentration, sugar content will also be enhanced under salinity. Consequently, soluble protein content (Fig. 5) was measured maximum in T11 (50% SBE) up to significant level (p<0.05) under induced salinity stress revealed that the soluble protein content has been promoted by increasing the concentrations of SBE. Nonetheless, total proline content (Fig. 6) of leaf was recorded highest in T11 by increasing SBE concentrations under induced salinity stress.

**Figure 4.**
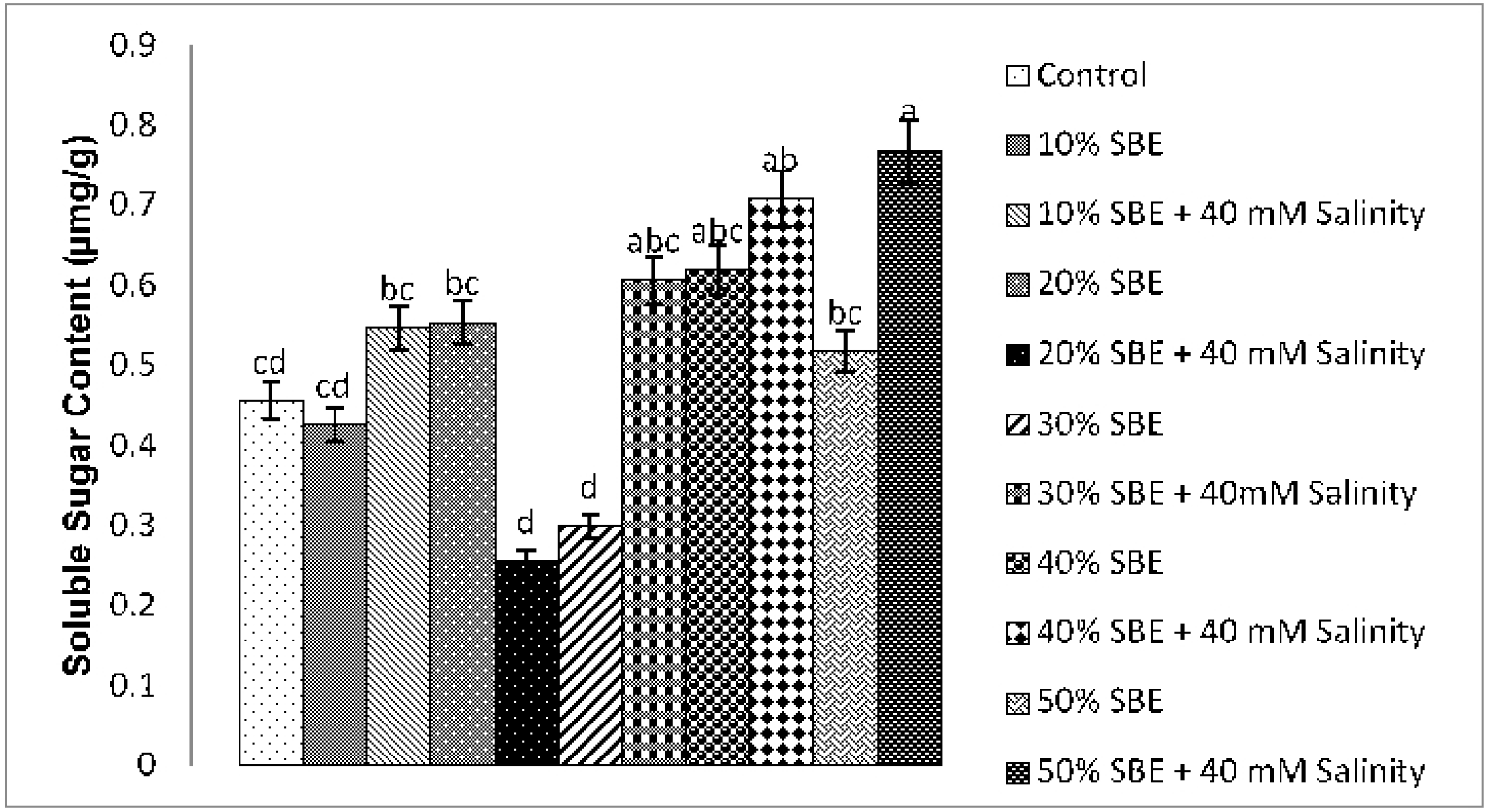
Effect of sugar beet extract on soluble sugar content under induced salinity stress. Vertical bars represent SE.

**Figure 5.**
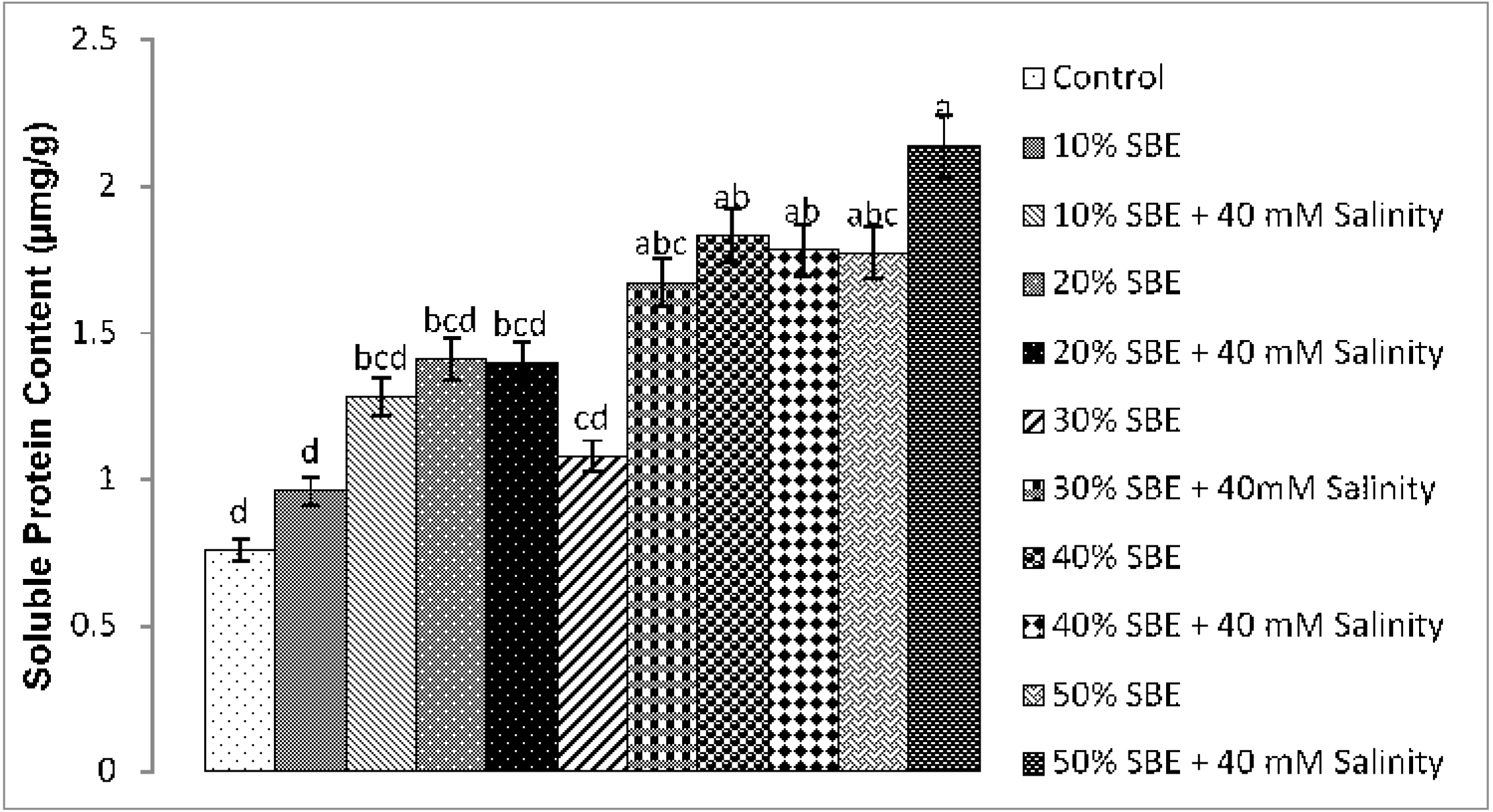
Effect of sugar beet extract on soluble protein content under induced salinity stress. Vertical bars represent SE.

**Figure 6.**
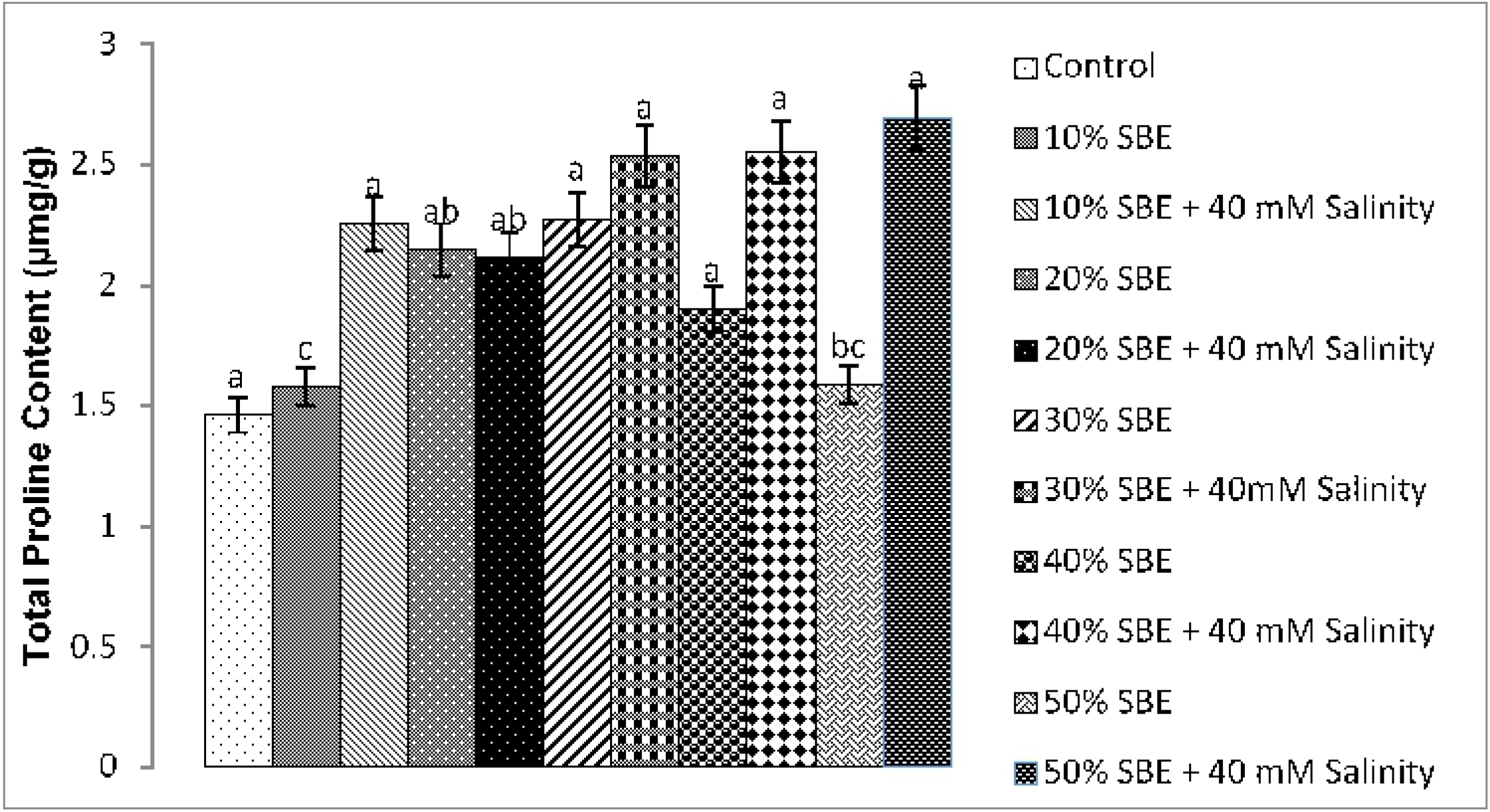
Effect of sugar beet extract on total proline contents under induced salinity stress. Vertical bars represent SE.

Formation of reactive oxygen species neutralize by various antioxidant enzymes by protecting the plants from harmful oxidative damage in stress environment is particularly unpredictable mechanism among different plant species and different cultivars of the same species (Hayat et al. 2012). The priming of seeds with different concentrations of SBE effectively reduced these enzymes activities by increasing sugar beet concentrations. Regarding SOD activity, 20% SBE concentrations are significantly more effective at improving SOD activity under salinity stress than other SBE levels (Fig. 7). However, in terms of POD activity, this improvement was greater in plants grown with 30% SBE primed under induced stress conditions (Fig. 8). Similarly, hydrogen peroxide activity studied in ‘Figure 9’ has been reported maximum in 30% sugar beet extract followed by 50% SBE. MDA is marker of lipid peroxidation/membrane loss under stress and indicates plant resistance to stress. In the current study, the MDA concentration augmented significantly enhanced under salinity in T11 stress as compared to plants grown under control and without salinity (Fig. 10).

**Figure 7.**
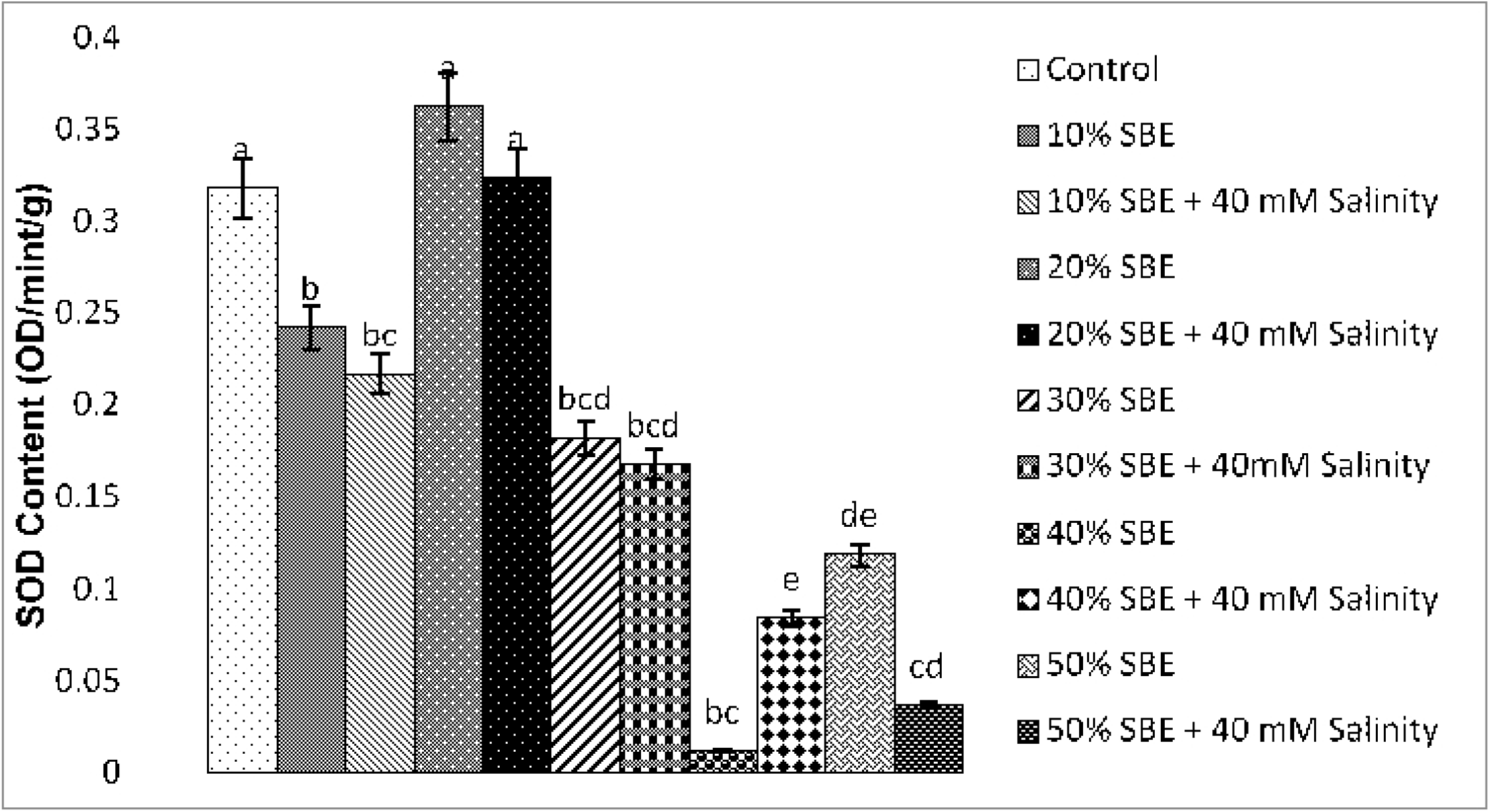
Effect of sugar beet extract on superoxide dismutase under induced salinity stress. Vertical bars represent SE.

**Figure 8.**
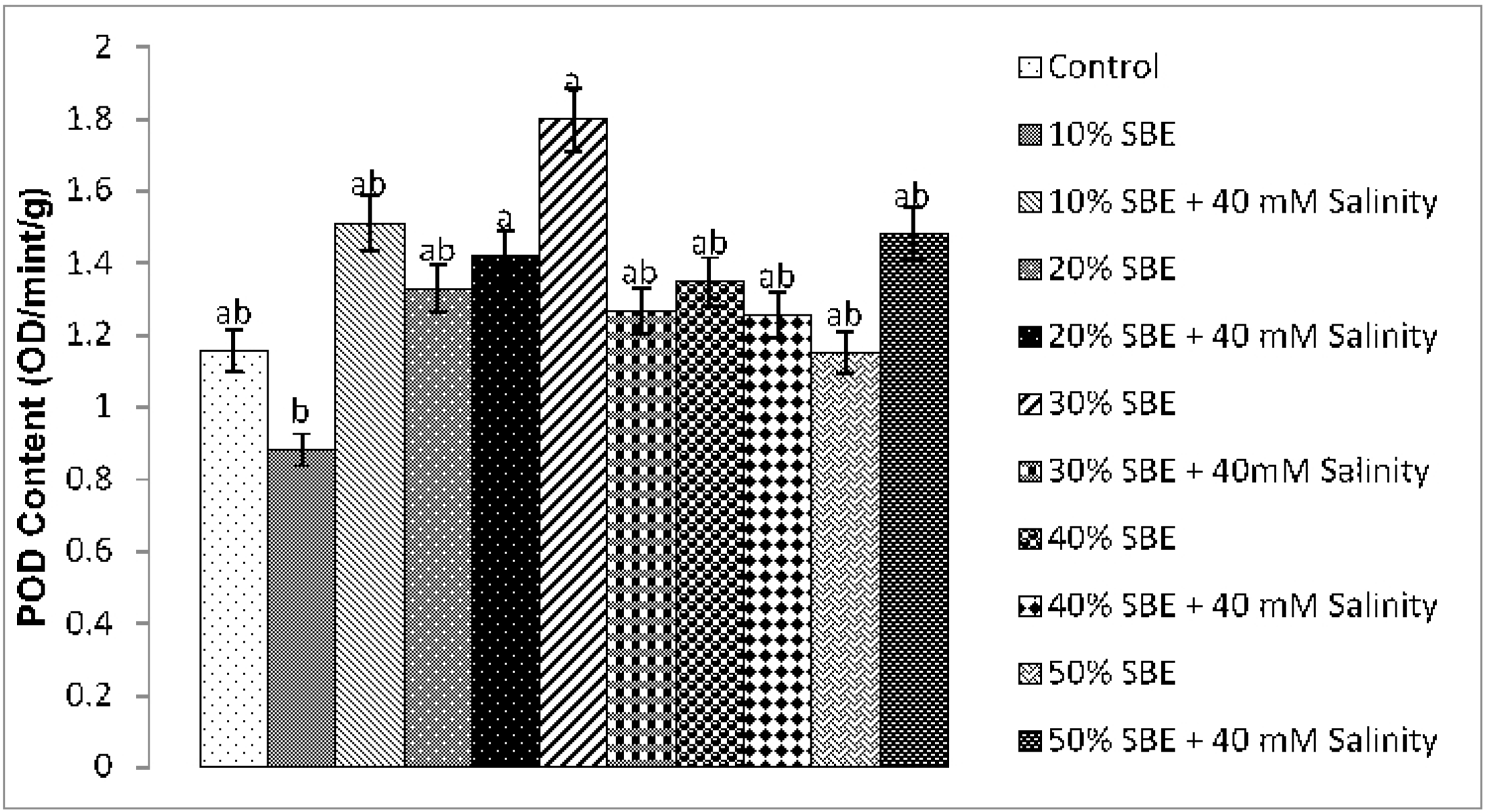
Effect of sugar beet extract on peroxidase under induced salinity stress. Vertical bars represent SE.

**Figure 9.**
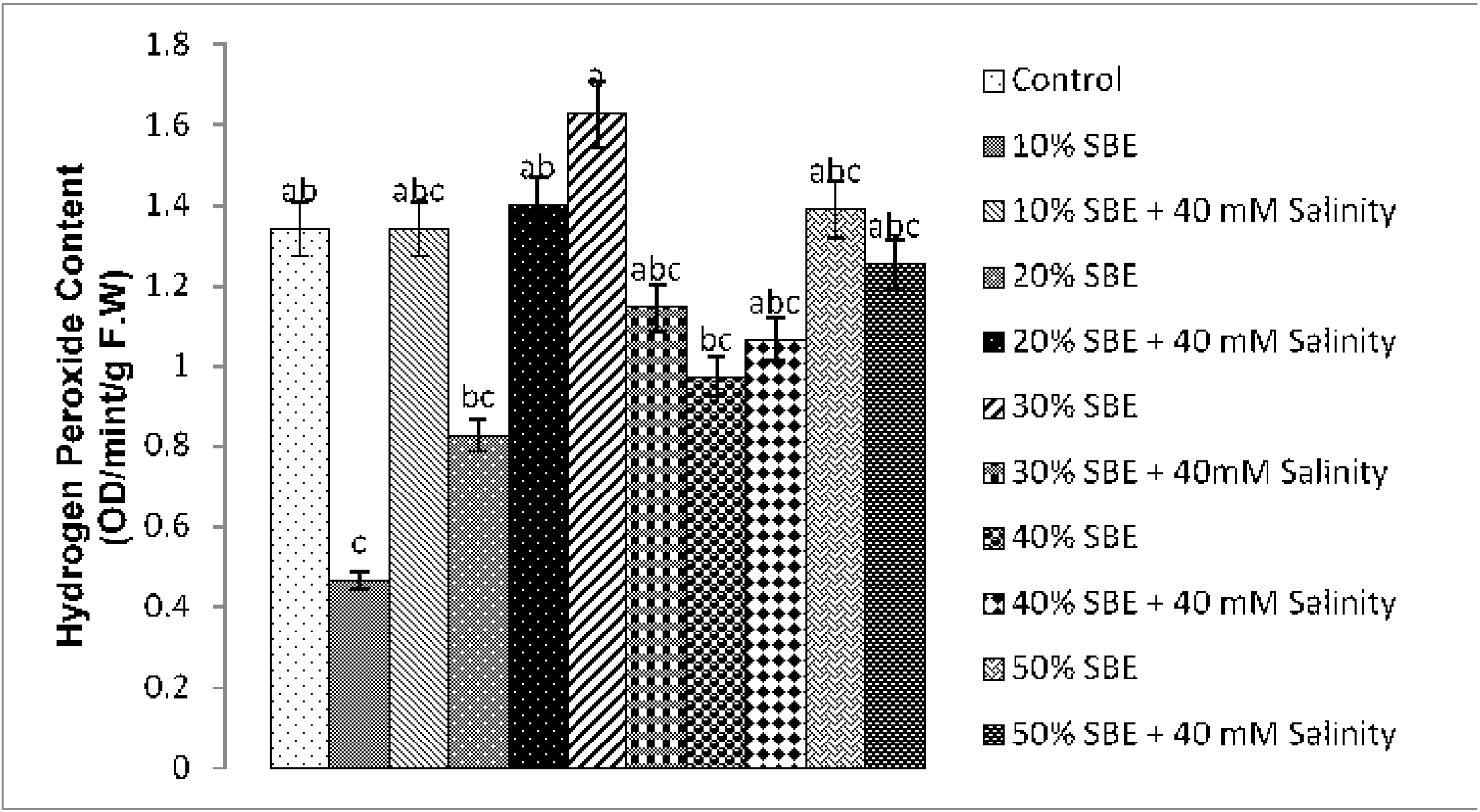
Effect of sugar beet extract on hydrogen peroxide under induced salinity stress. Vertical bars represent SE.

**Figure 10.**
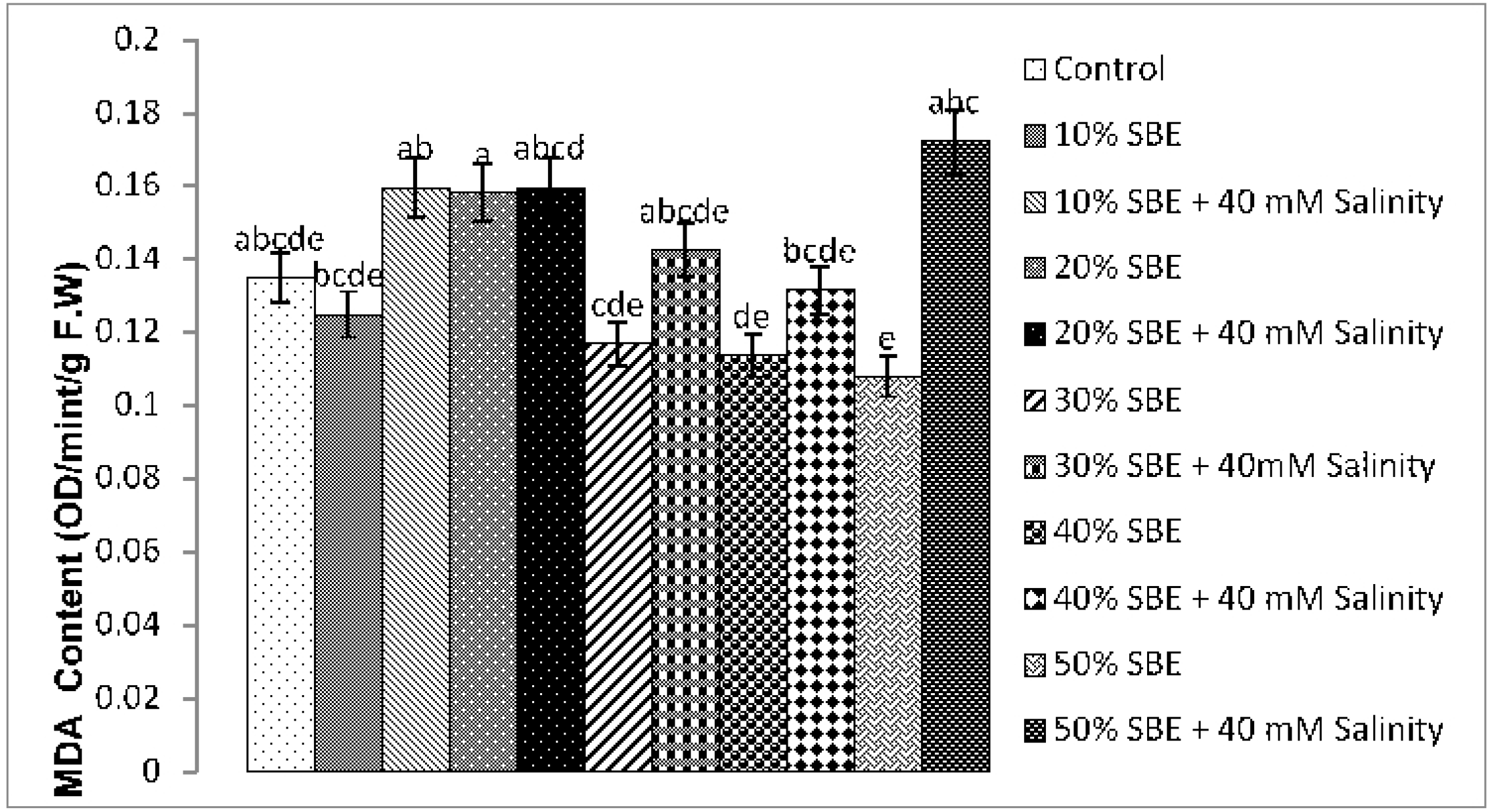
Effect of sugar beet extract on MDA content under induced salinity stress. Vertical bars represent SE.

### 3.3. Regression and correlation analysis of the measured traits

Statistical analysis showed that sugar beet extract affected on hydrogen peroxide activity in crop plants. Analysis of variance of measured traits with significant differences between treatments and within treatments was carried out between the priming factors under Salinity Stress (Table 4).

**Table 4.** Analysis of variance of measured traits under salinity stress in *Hordeum vulgare* L.

| **Traits** | **Source of variation** | **SS** | **Df** | **MS** | **F** | **P** |
| --- | --- | --- | --- | --- | --- | --- |
| **Chlo a/b ratio** | Between treatments | 68.647 | 10 | 6.865 | 0.771 | 0.655 |
|  | Within treatments | 195.985 | 22 | 8.908 | - | - |
|  | Error | 264.633 | 32 | - | - | - |
| **TCC** | Between treatments | 6.948 | 10 | 0.695 | 2.350 | 0.046* |
|  | Within treatments | 6.505 | 22 | 0.296 | - | - |
|  | Error | 13.453 | 32 | - | - | - |
| **CC** | Between treatments | 5.529 | 10 | 0.553 | 2.956 | 0.016* |
|  | Within treatments | 4.115 | 22 | 0.187 | - | - |
|  | Error | 9.644 | 32 | - | - | - |
| **SSC** | Between treatments | 0.742 | 10 | 0.074 | 4.855 | 0.001** |
|  | Within treatments | 0.336 | 22 | 0.015 | - | - |
|  | Error | 1.078 | 32 | - | - | - |
| **SPC** | Between treatments | 5.318 | 10 | 0.532 | 3.126 | 0.012* |
|  | Within treatments | 3.743 | 22 | 0.170 | - | - |
|  | Error | 9.062 | 32 | - | - | - |
| **TPC** | Between treatments | 5.450 | 10 | 0.545 | 3.872 | 0.004* |
|  | Within treatments | 3.097 | 22 | 0.141 | - | - |
|  | Error | 8.547 | 32 | - | - | - |
| **POD** | Between treatments | 1.662 | 10 | 0.166 | 0.851 | 0.588 |
|  | Within treatments | 4.296 | 22 | 0.195 | - | - |
|  | Error | 5.959 | 32 | - | - | - |
| **SOD** | Between treatments | 0.231 | 10 | 0.023 | 11.522 | 0.000*** |
|  | Within treatments | 0.044 | 22 | 0.002 | - | - |
|  | Error | 0.275 | 32 | - | - | - |
| **H<sub>2</sub>O<sub>2</sub>** | Between treatments | 4.493 | 10 | 0.449 | 1.494 | 0.207 |
|  | Within treatments | 6.615 | 22 | 0.301 | - | - |
|  | Error | 11.108 | 32 | - | - | - |
| **MDA** | Between treatments | 0.011 | 10 | 0.001 | 2.455 | 0.038* |
|  | Within treatments | 0.010 | 22 | 0.000 | - | - |
|  | Error | 0.022 | 32 | - | - | - |

Chlo a/b ratio=chlorophyll a/b ratio, TCC=total chlorophyll content, CC=caroteniod content, SCC=soluble sugar content, SPC=soluble protein content, TPC=total proline content, POD=peroxidase content, SOD=superoxide dismutase content, H<sub>2</sub>O<sub>2</sub>= hydrogen peroxide, MDA=malendyaldehyde, \* is significant up to p=0.01,\*\* is significant to p=0.01,\*\*\* is significant up to p=0.001.

According to the results of (Table 5, 6), regression and correlation analysis showed a significant positive correlation (p ≤0.05) between total chlorophyll and carotenoids content. Similarly, a positive correlation was also found between SSC, SOD, POD and MDA content in the leaves at p<0.05.

**Table 5.** Regression and correlation analysis between physiological and biochemical parameter in *Hordeum vulgare* L. under induced salinity stress.

| Chlo a/b | TCC | CC | SSC | SPC | TPC | POD | SOD | H<sub>2</sub>O<sub>2</sub> | MDA |
| --- | --- | --- | --- | --- | --- | --- | --- | --- | --- |
| R=0.034<sup>a</sup> | R=0.269<sup>a</sup> | R=0.211<sup>a</sup> | R=0.460<sup>a</sup> | R=0.701<sup>a</sup> | R=0.250<sup>a</sup> | R=0.122<sup>a</sup> | R=0.653<sup>a</sup> | R=0.090<sup>a</sup> | R=0.199<sup>a</sup> |
| R<sup>2</sup>=0.001 | R<sup>2</sup>=0.068 | R<sup>2</sup>=0.044 | R<sup>2</sup>=0.211 | R<sup>2</sup>=0.491 | R<sup>2</sup>=0.063 | R<sup>2</sup>=0.015 | R<sup>2</sup>=0.427 | R<sup>2</sup>=0.008 | R<sup>2</sup>=0.040 |
| Chlo a/b | TCC | CC | SSC | SPC | TPC | POD | SOD | H<sub>2</sub>O<sub>2</sub> | MDA |
| R=0.098 | R=0.167 | R=0.299<sup>a</sup> | R=0.055<sup>a</sup> | R=0.649<sup>a</sup> | R=0.450<sup>a</sup> | R=0.505<sup>a</sup> | R=0.461<sup>a</sup> | R=0.104<sup>a</sup> | R=0.058<sup>a</sup> |
| R<sup>2</sup>=0.010 | R<sup>2</sup>=0.174 | R<sup>2</sup>=0.090 | R<sup>2</sup>=0.003 | R<sup>2</sup>=0.421 | R<sup>2</sup>=0.065 | R<sup>2</sup>=0.203 | R<sup>2</sup>=0.213 | R<sup>2</sup>=0.011 | R<sup>2</sup>=0.003 |

Chlo a/b ratio=chlorophyll a/b ratio, TCC=total chlorophyll content, CC=carotenoid content, SSC=soluble sugar content, SPC=soluble protein content, TPC=total proline content, POD=peroxidase content, SOD=superoxide dismutase content, H<sub>2</sub>O<sub>2</sub>=hydrogen peroxide, MDA=malendyaldehyde, \* is significant up to p=0.01, \*\* is significant to p=0.01, \*\*\* is significant up to p=0.001.

**Table 6.**
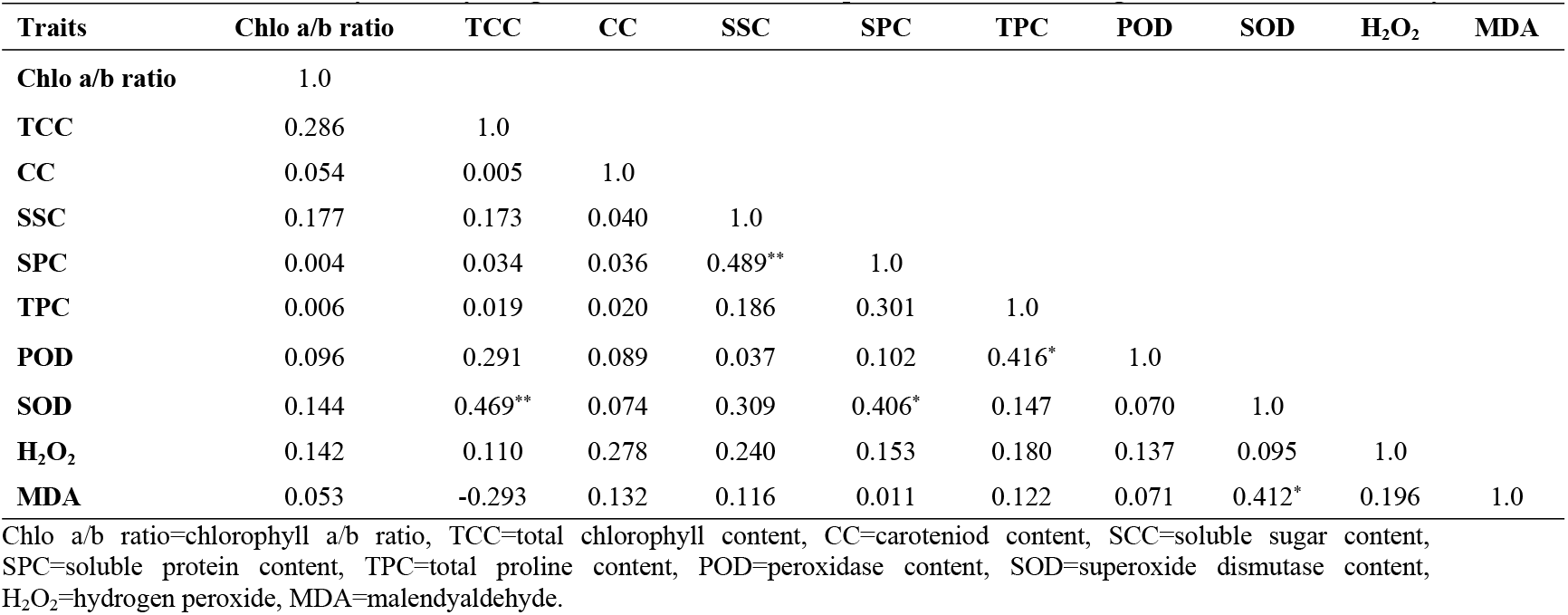
Multicorrelation Analysis of Physiological and Biochemical Components in *Hordeum vulgare* L. nder induced salinity stress.

### 3.4. Principle component analysis of biological component

Principle component analysis were studied on 10 characters (Table 7) enclosed over all 55.009% of the whole variation. The PC1 explained 23.562% of complete variance which were significantly correlate with protein, proline, soluble sugar content and carotenoid content particularly related with osmolytes. while PC2 investigated 17.694% of complete variance which was particularly correlate with total chlorophyll content, chlorophyll “a/b” ratio, SOD and POD corresponded to plant antioxidant enzymes and growth components.

**Table 7.** Principal component based on correlation matrix of biological component in *Hordeum vulgare* L.

| Components | Eigen value | Individual | Cumulative | PC1 | PC2 | PC3 |
| --- | --- | --- | --- | --- | --- | --- |
| **Chlo a/b ratio** | 1.788 | 13.753 | 55.009 | 0.337 | 0.249 | 0.496 |
| **TC** | 1.306 | 10.050 | 65.059 | 0.921 | 0.235 | 0.140 |
| **CC** | 1.111 | 8.547 | 73.605 | 0.080 | 0.102 | 0.425 |
| **SSC** | 1.000 | 7.690 | 81.296 | 0.313 | 0.596 | 0.390 |
| **SPC** | 0.725 | 5.574 | 86.870 | 0.280 | 0.742 | 0.055 |
| **TPC** | 0.582 | 4.480 | 91.350 | 0.067 | 0.563 | 0.282 |
| **POD** | 0.460 | 3.537 | 94.887 | 0.192 | 0.396 | 0.612 |
| **SOD** | 0.330 | 2.535 | 97.422 | 0.733 | 0.337 | 0.125 |
| **H<sub>2</sub>O<sub>2</sub>** | 0.203 | 1.565 | 98.987 | 0.158 | 0.164 | 0.640 |
| **MDA** | 0.132 | 1.013 | 100.000 | 0.422 | 0.283 | 0.236 |

Chlo a/b ratio= chlorophyll a/b ratio, TCC= total chlorophyll content, CC = carotenoid content, SCC= soluble sugar content, SPC = soluble protein content, TPC= total proline content, POD= peroxidase content, SOD = superoxide dismutase content, H<sub>2</sub>O<sub>2</sub>= hydrogen peroxide, MDA= malendyaldehyde.

## 4. Discussion

Irrespective of hundreds of factors causing impediments in plant growth and development, water shortage is reckoned as a main abiotic factor that affects the crop growth and limits the global food production (Noman et al. 2018). Sugar beet extract is a vital supply of flavonoids, glycine betaine, betalains, phenolics, ascorbic acid (AsA) etc. and is found to be most effective as a bio-stimulant described by the ECBI (European Council of Bio-stimulant Industry) (Colla et al. 2015; Nyoni et al. 2020). The current study revealed that priming with SBE enhanced germination energy (GE), means germination time (MGT), coefficient of velocity of emergence (CVE) as compared to control. Our results are according to the findings of Noman et al. (2020) who reported significant enhancement in GE, MET and CVE by priming seeds with SBE under water stressed wheat. Among the treatments of 10% SBE and 30% SBE are more effective to salinity stress. This suggested that sugar beet extract could be better treatment of seed priming for crops (Ansari and Zede 2012). Similarly, salt stress limited the levels of C, N and O in barley leaves. It has been suggested that pre-sowing seed treatment with concentrated SBE endeavored to balance the reduced minerals in plants under salinity stress as well as germination parameters. (Abbas et al. 2010; Liu et al. 2020).

Differences in photosynthetic pigments are a key indicator for determining the photosynthetic pigments of plants under salinity stress. Our results (Fig. 1-3) showed a noticeable increase in chlorophyll “a/b ratio, total chlorophyll and carotenoids by pre-sowing seeds with SBE both with and without salinity stress. Our findings are correlated with the study of Du-Jardin (2015), Ali and Ashraf (2011) who suggested that SBE act as a natural biostimulant against salinity stress. However, carotenoids content increased under salinity due to SBE seed priming might have a defensive role by reducing the damages to photosystem II proteins due to reduced light harvesting.

The osmolytes in leaf such as sugar and proline level by priming with SBE were measured highest under induces salinity stress that proved to be effective by the adverse effect of stress condition in barley. Similar results have been found by Abbas et al. (2010) who studied that proline concentration in the leaves of both eggplant cultivars increased markedly with salt treatment. Furthermore, it is estimated that different concentrations of sugar beet extract enhanced the proline contents under induced salinity stress, respectively (Abdelaal et al. 2020; Zeeshan et al. 2020). Similarly increase in soluble protein under salinity stress via seed priming with 50% SBE may be due to broadspectrum of stress resistance that has been unlocked by osmolytes to survive under several biotic and abiotic stresses. Our results are corelated with the finding of Noman et el. (2018) who studied that SBE seed pre-sowing further improved the total protein content of plants grown under both control and stressed conditions.

To reduce the negative effects of ROS, plants have well-developed antioxidant defense mechanisms that include the activity of well-known antioxidant enzymes including CAT, POD, SOD, and APX (Zeeshan et al. 2020). Antioxidant enzymes such as POD, SOD, MDA activities enhanced by pre-sowing seeds with SBE while H_2_O_2_ formation was maximum under applied salinity stress. Increased activity of SOD indicates the efficient destruction of superoxide ion to H_2_O_2_ which is eliminated by POD (Mollern 2007) whereas, GB (glycine betaine) present in SBE enhanced osmoprotectant level which reduce ROS formation by acting as antioxidative compound (Jamil et al. 2015). Similar results have been indicated by *Ebrahim* et al. (2020) who reported that the sugar beet extract was significant under the induced salinity stress.

## 5. Conclusion

The present study investigated that seed priming with SBE is a reliable and cost-effective technique for the agricultural and seed quality. It can be evaluated that seed priming treatment with sugar beet extract is of significant value for barley plants facing osmotic stress. A clear correspondence between modulated concentrations of mineral ions, antioxidative enzymes and germination indices was measured in barley plants elevated from seeds treated with SBE under salinity stress. Overall, 30, 40 and 50% concentration SBE exerted a positive effect on germination, physiobiochemical under both untreated primed as well as saline treated stressed condition. Nevertheless, further research on SBE will be useful for seed quality and germination parameters as still there is a demand for additional exploration of plant stress tolerance mechanism by priming seeds with SBE through involving modeling and experimental approaches. Forthcoming findings would be performed utilizing varied cultivars/lines for drawing parallels among SBE application under field conditions, salinity stress, ionic mobility patterns, crop quality and production.

## Abbreviations

SBE: sugar beet extract
CVE: coefficient of velocity of emergence
MET: mean emergence time
GE: germination energy
TGI: timson germination index
GRI: germination rate index
POD: Peroxidase
MDA: Malondialdehyde
SOD: Superoxide dismutase
H_2_O_2_: Hydrogen peroxide
ROS: reactive oxygen species
AsA: Ascorbic acid
CAT: Catalase
TSS: total soluble sugars (TSS)
G1: Germinated seed percentage at day first

## Acknowledgements

We are highly acknowledged to Department of Botany, University of Peshawar for providing all facilities regarding this work.

## Data Availability

Data associated with the present paper can be obtained by contacting the corresponding authors.

## Authors’ contributions

Sami Ullah designed the experiments and supervised the experiments of the first author Noor Ali Shah who conducted the experiments, wrote manuscript and analysed all data. Sajjad Ali, Muhammad Adnan and Muhammad Nauman Khan conduct antioxidants and biochemical analysis, Amjad Ali, Ajmal Khan and Said Hassan did statistical analysis and help in manuscript writing, Muhammad Ali and Wisal Muhammad Khan reviewed and checked the final version. All authors have read and agreed to its contents and also that the manuscript complies with the policy of the publication.

